# Purinergic P2X7 Receptor-mediated inflammation precedes PTSD-related Behaviors in Rats

**DOI:** 10.1101/2022.03.10.483788

**Authors:** Orlando Torres-Rodriguez, Yesenia Rivera-Escobales, Bethzaly Velazquez, María Colón, James T. Porter

**Author notes:** Correspondence should be addressed to: James T. Porter, Ph.D., Ponce Health Sciences University-Ponce Research Institute, PO Box 7004 Ponce, PR 00732-7004.

## Abstract

Clinical evidence has linked increased peripheral pro-inflammatory cytokines with post-traumatic stress disorder (PTSD) symptoms. However, whether inflammation contributes to or is a consequence of PTSD is still unclear. Previous research shows that stress can activate P2X7 receptors (P2X7Rs) on microglia to induce inflammation and behavioral changes. In this investigation, we examined whether P2X7Rs contribute to the development of PTSD-like behaviors induced by single prolonged stress (SPS) exposure in rats. Consistent with the literature, exposing adult male and female rats to SPS produced a PTSD-like phenotype of impaired fear extinction and increased anxiety-like behavior one week after exposure. In addition, SPS-exposed animals had more Iba1-positive microglia expressing the P2X7R in the ventral hippocampus, a structure that regulates fear extinction and anxiety-like behavior. Next, we examined if inflammation precedes the behavioral manifestations. Three days after SPS exposure, increased inflammatory cytokines were found in the blood and hippocampal microglia showed increased expression of the P2X7R, IL-1β, and TNF-α, suggesting increased peripheral and central inflammation before behavioral testing. To determine whether P2X7Rs contribute to the PTSD-related behaviors induced by SPS exposure, we gave ICV infusions of the P2X7R antagonist, A-438079, for one week starting the day of SPS exposure. Blocking P2X7Rs prevented the SPS-induced impaired fear extinction and increased anxiety-like behaviors in male and female rats, suggesting that SPS activates P2X7Rs which increase inflammation to produce a PTSD-like phenotype.

## Introduction

Post-traumatic stress disorder (PTSD) is a debilitating neuropsychiatric disorder that can develop following exposure to a traumatic incident [1]. Recent reviews found increased levels of IL-6, IL-1β, TNF-α, and IFNƔ in the serum of PTSD patients suggesting an inflammatory signature as a potential biomarker for PTSD [2,3]. Likewise, a positive correlation was found between peripheral IL-1β, IL-6, and TNF-α and PTSD severity [4]. Although these reports suggest that inflammation is associated with PTSD, whether the inflammation is causal in the PTSD pathophysiology is unclear.

A previous study found that activation of the innate immune response by LPS impairs fear extinction in rats [5]. This study implicates the immune system as a potential contributor to impaired fear extinction learning. In addition, exposure to single prolonged stress (SPS) induces the activation of microglia and increases hippocampal IL-1β and TNF-α [6]. Furthermore, SPS exposure increases inflammatory responses in the basolateral amygdala (BLA) [7], hippocampus [6,8], and medial prefrontal cortex [8]. Since SPS is a well-established preclinical model that induces PTSD-related behaviors in rodents [9,10], stress-induced inflammation from microglia might be triggering PTSD-related behaviors such as the impaired fear extinction observed seven days after SPS in male rodents [8,11]. Although the literature suggests a close relationship between stress-induced inflammation and PTSD-related behaviors, whether inflammation precedes and contributes to the induction of PTSD-related behaviors is unclear.

The release of ATP during stress [12] could stimulate P2X7Rs, expressed predominately by microglia [13,14], to cause a central inflammatory response to induce the PTSD-related impaired fear extinction. To test this hypothesis, we examined the expression of P2X7Rs and inflammatory cytokines by hippocampal microglia three days after SPS which is before the development of impaired fear extinction [8,11], and found increased cytokine and P2X7R expression. Furthermore, we found that pharmacological inhibition of P2X7Rs during SPS prevented the development of impaired auditory fear extinction and increased anxiety-like behavior one week later indicating that SPS induces PTSD-related behaviors through P2X7R activation.

## Materials &Methods

### Animal subjects

All animal procedures were approved by the Institutional Animal Care and Use Committee of the Ponce Health Sciences University (PHSU) in compliance with NIH guidelines for the care and use of laboratory animals. Adult male and female Sprague Dawley rats were transported from the PHSU colony to a satellite facility nearby where they were individually housed on a 12/12 h light/dark schedule with free access to food and water.

### SPS

Sprague-Dawley rats approximately post-natal day 60 were pseudorandomly assigned to the SPS or the non-stressed (NS) group. As originally described [9,11], SPS started with 2 hours of restraint stress using a disposable rodent restrainer (DecapiCone®; Cat. No. DC-200), followed by immediate exposure to a 20-minute forced swim in a cylinder (20cm X 45cm) containing tap water at 24°C. Next, rats recovered from the physical stress in a cage under direct light (soft white 60 watts bulb) as a heat source for 10-min. Then, we placed the rats in an anesthetic chamber (16cm X 16cm) with ethyl ether (Millipore Corporation, Cat. No. EX0185-8) until general anesthesia induction. Each animal received SPS individually. Following SPS, animals were single-housed and left undisturbed for seven days before behavioral testing. The NS group was housed under identical conditions.

### Differential Auditory Fear Conditioning (DAFC)

NS and SPS groups were exposed to DAFC and extinction to test their ability to discriminate between a safe and an aversive cue. The 3-day fear discrimination phase consisted of two auditory cues; a safe conditioned stimulus (CS-, 7 kHz, 80 dB) and an aversive conditioned stimulus (CS+, 1 kHz, 80 dB) paired with a footshock. During day 1, animals received six CS-tones with a 3-minute Intertrial Interval (ITI) in context A. During day 2, animals received six CS+ tones, one habitation tone and five tones paired with a 0.44 μA footshock with a 3-minute ITI in context A. During the discrimination test on day 3, rats received two CS-and two CS+ tones in a novel context B with an hour between sessions. Context A was a clear acrylic cage with electrified grid floor (ID#46002; Ugo Basile). In context B the visual, tactile, and olfactory cues were changed to reduce contextual effects on the cued responses.

### Fear Extinction

Following the discrimination test, animals received two fear extinction sessions in a novel context B on day 4. Each extinction session consisted of 14 CS+ tones with a 3-minute ITI in context B. The animals were returned to their home cage undisturbed for one hour in between extinction sessions. On day 5, animals received two CS+ tones to test their fear extinction memory.

### Anxiety-like Testing

On day 6, we exposed the animals to a 20-minute open field test (OFT) in a 94cm X 94cm X 44cm arena to assess anxiety-like behavior. During the OFT, a total of 6 CS+ tones were presented to assess the cue-associated anxiety-like behavior of the animals.

### Enzyme-linked immunosorbent assay (ELISA)

Following behavioral testing, both groups were deeply anesthetized and sacrificed for sample collection. Trunk blood was collected using the BD Vacutainer ® K2 EDTA (K2E) 3.6 mg blood collection tubes (Cat. No. 367841), centrifuged at 3,000 G for 5 minutes, and the supernatant (plasma) was stored at -80°C until further processing. Plasma samples were diluted 1:10 in the 1X diluent buffer provided by the Rat Cytokine ELISA plate kit (Signosis, Cat. No. EA-4006) for the analysis of 16 inflammatory cytokines using the Rat Cytokine ELISA plate (Signosis). Absorbance values at 450nm detected by the MultiSkan Go (Thermo Scientific) were used to determine relative cytokine expression.

### Hippocampal Microglial Isolation and RNA Extraction

Three days after SPS, we removed the brain and dissected the whole hippocampus from both hemispheres for microglial isolation. We enzymatically and mechanically dissociated hippocampal tissue as indicated in the Adult Brain Dissociation Kit (Miltenyi Biotec; Cat. No. 130-107-677). Then, we eliminated the cell debris and red blood cells using the same kit. We incubated the cell suspension with 5 ul of CD-11β/c Microglia MicroBeads (magnetic beads Miltenyi Biotec; Cat. No. 130-105-643). Next, we passed the cell suspension through the MACS columns (Miltenyi Biotec; Cat. No. 130-105-634) for microglial isolation. We immediately extracted RNA from the isolated CD-11β+ cells as instructed by the Arcturus® picoPure® RNA isolation Kit (Applied Biosystems Cat. No. 12204-01). The microglial RNA portion for each sample was approximately 30 μl and was stored at -80°C until further processing.

### Real-time Polymerase Chain Reaction (RT-PCR)

Purified microglial RNA was processed for cDNA synthesis using the iScript cDNA synthesis kit (BIO-RAD, 1708891). The cDNA was diluted 1:20 using Molecular Biology Reagent Water (Sigma Life Science, Cat No. W4502-1L). RT-PCR was performed using primers for P2X7R, IL-1β, CD68, TNF-alpha, and iQ SYBR Green Supermix (BIO-RAD, Cat. No. 1708882). Cycle threshold values were normalized to housekeeping gene GAPDH (Integrated DNA Technologies). Each experiment was performed in duplicate and values were averaged.

### Immunofluorescence Staining

We collected brain samples 24 hours after behavioral testing. Brain samples were fixed, dehydrated, and embedded in paraffin. We mounted paraffin-embedded VH coronal slices (4 μm) onto positively charged slides. Tissue was deparaffinized in xylene and rehydrated in a descending CDA19 ethanol series. Antigen retrieval consisted of incubation with 0.01M Citrate-EDTA solution (pH = 6.2) for 40 minutes followed by a 20-minute incubation at room temperature. The slides were incubated overnight in a humidified chamber at 4°C with the primary antibodies for Iba1 (1:1,000; Wako Chemicals; Cat. No. 019-19741) and P2X7R (1:200; Santa Cruz Biotechnologies; Cat. No. sc-134224). Primary antibodies were labeled with Alexa Fluor 568 Goat Anti-Rat (Cat. No. A-11077) for P2X7R and Alexa Fluor 488 Goat Anti-Rabbit (Cat. No. A-21206) for Iba1. A control reaction was performed without primary antibodies for each test. Tissues were covered with ProLong™ Gold antifade reagent (Thermo Fischer Scientific; Cat. No. P36934) and a cover slide. Nuclei were stained with NucBlue Fixed Cell Stain (DAPI, Cat. No. 12333553). Images were taken using an Olympus Microscope (Model BX60), Nikon digital camera (Nikon, DS-Fi1), and camera control unit (Nikon, DS-U2) with the NIS Elements software (AR 2.22.15). Images of tissues from the P2X7R antagonist experiment were taken using a Nikon Confocal Microscope A1 (Ver.4.10). Images were acquired by investigators blinded to treatment groups. Cell counting and fluorescence analyses of all images were performed using the cell-counter plug-in of the ImageJ software (NIH, USA) by investigators blinded to treatment groups. Only cells with clear DAPI-stained nuclei were counted as cells.

### Stereotaxic Surgery and intracerebroventricular (ICV) administration of P2X7R antagonist

Two weeks before SPS exposure, animals were placed in a stereotaxic frame, anesthetized with 2%-2.5% isoflurane and oxygen, and implanted with a cannula (7 mm) in the left cerebral ventricle (AP, - 0.8; ML,1.4; DV, 3.8mm) that was fixed in place. Animals recovered for two weeks before behavioral training (Figure 1A). A microsyringe delivered ICV infusions of A-438079 (Tocris; Cat. No. 2972) or vehicle. Each animal received a total of 2 μl at a rate of 0.5 μl/min of either VEH (0.9% NaCl solution) or (4.47 mM A-438079). We selected the dose based on reported efficacy [15]. Following the behavioural procedures, animals were given a lethal dose of Euthanasia-III Solution (Pentobarbital Sodium, Phenytoin Sodium, MED-PHARMEX™) and the brain was dissected for molecular examination.

**Figure 1:**
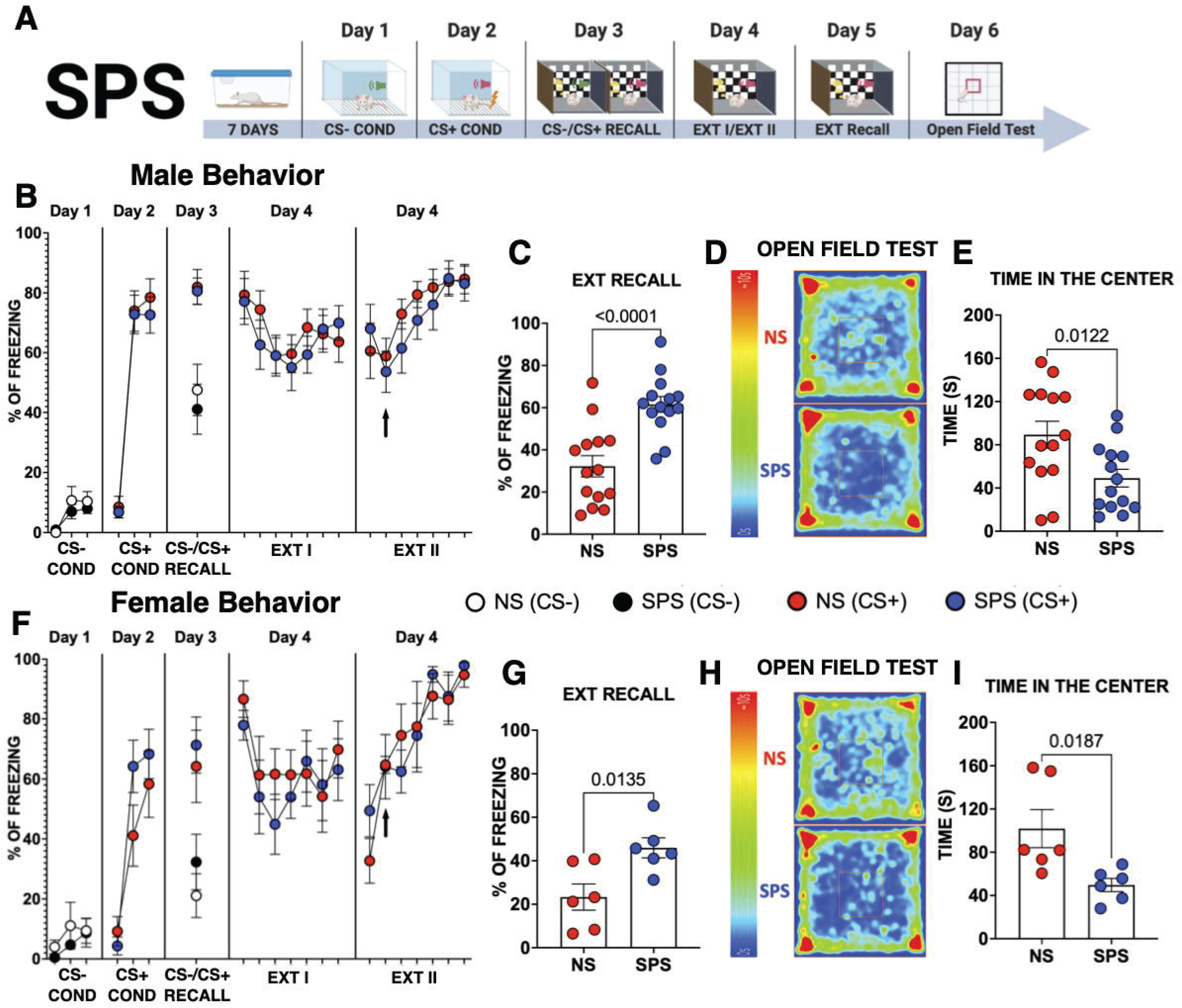
SPS impairs fear extinction recall and increases anxiety-like behaviors. (A) Representation of the Differential Auditory Fear Conditioning (DAFC) paradigm. (B, F) Graph of the percent freezing of male (B) and female (F) rats across the behavioral paradigm. (C, G) Percentage of freezing during EXT-Recall on Day 5 for male (C) and female (G) groups. (D, H) Average heat maps of the center of the body for male (D) and female (H) groups. (E, I) Graphs of the time spent in the center of the OFT for males (E) and female (I) rats.

### Data analysis

All behavioral and molecular data were analyzed with Graphpad Prism (version 9.1.0, San Diego, California). Auditory fear was measured as the percent of time spent freezing during each 30 second tone of training and recall with ANY-maze software (Ugo Basile, Italy). The two trials of fear recall were averaged. Data are presented as the mean ± SEM. Unpaired t-tests were utilized for group comparisons (Graphpad Prism version 9.1.0, San Diego, California). Two-way analysis of variance (ANOVA) for repeated measures were employed for comparisons between treatment groups over time. Sidak’s Multiple Comparisons were used for post hoc comparisons when appropriate. Significance was set at p ≤ 0.05.

## Results

### SPS impaired fear extinction and increased anxiety-like behaviors

First, we examined whether SPS exposure disrupts the cued-associated fear discrimination by exposing the animals to a DAFC training and testing (Figure 1A). As expected, both NS and SPS animals did not present fear responses (freezing) to the CS-tone (Figure 1B, F, Day 1). The percent freezing to the last CS-tone was not different between NS and SPS males (t (26) = 0.5338, p = 0.5980) and female groups (t (10) = 0.8474, p = 0.3658). The next day, all rats received five CS+ tones paired with a footshock. Both NS and SPS groups associated the CS+ tone with an aversive outcome (Figure 1B-F, Day 2). All groups froze more to the last CS+ than the first CS+ (Statistical Table). In addition, the NS and SPS male (t (26) = 0.6821, p = 0.5012) and female (t (10) = 0.7167, p = 0.4899) groups showed similar fear responses to the last CS+ during training on day 2, suggesting that SPS did not enhance cued fear learning in either sex. On day 3, we tested whether SPS impairs the specificity of the cued fear learning by measuring fear discrimination. NS and SPS exposed animals exhibited less freezing to the CS-than to the CS+ in males (t (26) = 3.436, p = 0.0020, NS; t (26) = 3.997, p = 0.0005, SPS) and females (t (10) = 3.613, p = 0.0047, NS; t (10) = 2.967, p = 0.0141, SPS). These results suggest that SPS did not disrupt cued fear discrimination in either sex. In addition, the freezing to the CS+ was similar in both the NS and SPS male (t (26) = 0.1875, p = 0.8527) and female (t (10) = 0.04487, p = 0.9651) groups suggesting that SPS did not enhance consolidation of the auditory fear memory.

The following day (Day 4), all groups received two sessions of the CS+ tones in a novel context B to examine the effect of SPS on fear extinction learning (Figure 1B, F). SPS did not alter cued fear extinction learning in males or females (Statistical table). Previous research reported that SPS impairs contextual and cued fear extinction memory [11] and increases anxiety [7,8] one week later in male rats. Consistent with the literature, we found that SPS-exposed male rats froze more during extinction recall than NS males (t (26) = 4.670, p = 0.0001) indicating that SPS impaired fear extinction memory in the males (Figure 1C). The next day, SPS-exposed male rats also spent less time in the center of the OFT (t (26) = 2.694, p = 0.0122) suggesting that SPS increased anxiety-like behavior in the male rats (Figure 1D, E). We also explored the effects of SPS on extinction recall and anxiety-like behaviors in female rats (Figure 1G-I). In contrast to a previous report [16], we found that SPS-exposed female rats also exhibited impaired extinction recall (t (10) = 2.993, p = 0.0135) and increased anxiety-like behavior (t (10) = 2.802, p = 0.0187) compared to NS female rats. Therefore, SPS impaired cued fear extinction recall and increased anxiety-like behaviors in both sexes.

### Increased ventral hippocampal expression of Iba1 and P2X7R in rats exposed to SPS

Animal and clinical studies suggest a close relationship between psychiatric illnesses and microglial activation [17–20]. Therefore, we examined whether SPS increased the expression of two inflammatory-associated microglial markers, Iba1 [21] and the P2X7R [13,14]. Although many different brain structures are involved in fear extinction and anxiety-like behaviors, we chose to examine the ventral hippocampus (VH) since it is central to both behaviors [22–24]. We collected VH brain slices and used immunofluorescence to examine Iba1 and P2X7R expression one day after behavioral testing. SPS-exposed male animals exhibited more Iba1 (t (11), 4.064, p = 0.0019) and P2X7R (t (11), 3.129, p = 0.0096) positive cells in the VH (Figure 2A-F). To verify that the cells expressing the P2X7R were microglia [14], we quantified the number of Iba1+/P2X7R+ cells in the VH (Figure 2G). We found that SPS-exposed male rats exhibited more Iba1+/P2X7R+ cells (t (11), 2.998, p = 0.0121) in the VH. Analysis of the percent area as a relative measure of protein expression was performed (Figure 2H-I). We found no difference in the expression of Iba1 (t (11) = 1.680, p = 0.1211), however SPS-exposed rats exhibited a higher percent area of P2X7R (t (10), 2.590, p = 0.0270). The increased expression of inflammatory markers suggests that exposure to SPS induced a pro-inflammatory state in the brain. However, this inflammatory manifestation could require both SPS and behavioral training.

**Figure 2:**
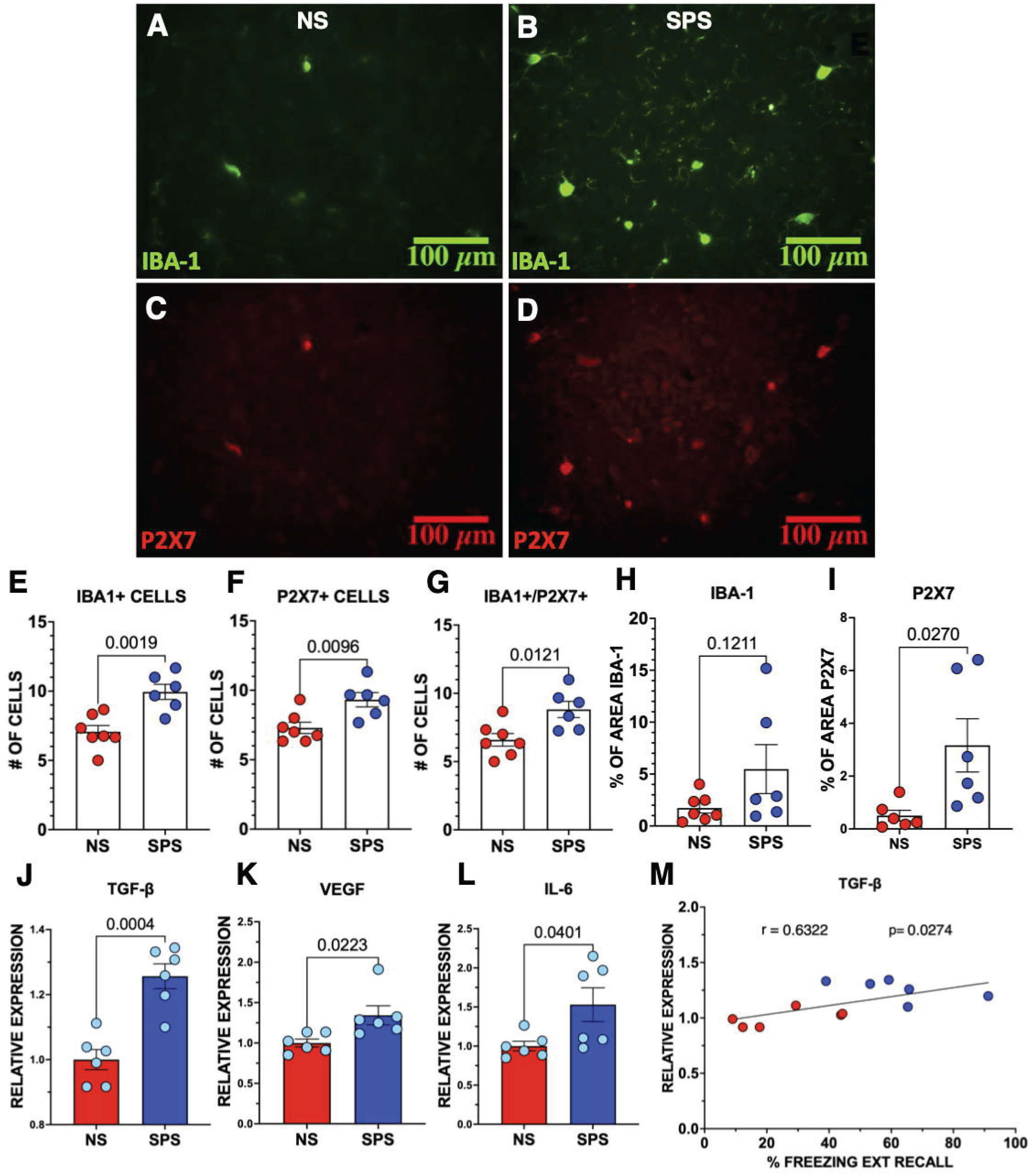
SPS increases VH Iba1 and P2X7R expressing cells and peripheral production of cytokines in male rats. (A, C) Representative images of VH slices from NS male rats co-labeled for (A) Iba1 and (C) P2X7R. (B, D) Representative images of the SPS-exposed male rats. (E-G) Quantification of Iba1+, P2X7R+, and Iba1+/P2X7R+ cells in the VH. (H-I) Graphs of area fraction of Iba1 and P2X7R immunofluorescence in VH slices. (J-L) Peripheral pro-inflammatory cytokines TGF-β, VEGF, and IL-6. Absorbance values were normalized to NS male group. (M) Correlation of peripheral TGF-β and percent freezing during EXT recall.

### SPS exposure increased peripheral pro-inflammatory cytokines

At the end of behavioral testing, all animals were sacrificed, and blood and brain tissue were collected for further analysis. Since increases in inflammatory cytokines have been found in the blood of patients with PTSD [3,25], we examined whether SPS altered the expression of 16 pro-inflammatory cytokines in blood plasma by ELISA. Consistent with clinical evidence suggesting that stress exposure is associated with peripheral production of inflammatory cytokines [25], the SPS-exposed male rats exhibited higher expression of several cytokines (Figure 2J-L) including TGF-β (t (10) = 5.209, p = 0.0004), VEGF (t (10) = 2.701, p = 0.0223), and IL-6 (t (10) = 2.358, p = 0.0401) with a trend towards an increase in IL-1β (t (10) = 1.982 p = 0.0756) and leptin expression (t (10) = 2.068, p = 0.0655). In contrast, the pro-inflammatory cytokine TNF-α (t (10) = 1.237, p = 0.2444) and several other cytokines were not increased (Supplemental Table 1). Of the three cytokines increased by SPS, only TGF-β positively correlated with the percentage of freezing during extinction recall (Figure 2M). These results suggest that SPS exposure increases the production of certain peripheral pro-inflammatory cytokines which could also contribute to the PTSD-related behaviors seen in the SPS-exposed rats.

### SPS increased hippocampal microglia pro-inflammatory genes 3 days post-exposure

The above results demonstrate that animals that already exhibit a PTSD-like phenotype also show signs of peripheral inflammation and microglial activation. These results raised the question of whether the SPS-exposed animals entered the behavioral paradigm with an increased inflammatory profile that contributes to PTSD-related behaviors. Since SPS-exposed animals exhibited more P2X7R+ microglia at the end of behavioral analysis, we examined whether SPS increased hippocampal microglial expression of the P2X7R gene prior to behavioral training. A previous report found that SPS increases protein expression of the high mobility group box 1 (HMGB1) in the BLA three days after SPS [7]. Therefore, we isolated hippocampal microglial RNA from male and female rats three days after SPS to determine whether the SPS-exposed animals enter the behavioral paradigm with an increased inflammatory profile. Since males and females showed similar behaviors, we combined both sexes for the molecular analysis. We found that hippocampal microglia from SPS-exposed animals expressed more P2X7R (t (34), 2.532, p = 0.0161), TNF-α (t (34), 2.673, p = 0.0115), IL-1β (t (34), 2.335, p = 0.0256), and CD68 (t (34), 2.641, p = 0.0124) 3 days after SPS (Figure 3A-D). These results suggest that the SPS-exposed animals entered the fear learning paradigm with an increased inflammatory profile which could contribute to the observed PTSD-related behaviors.

**Figure 3:**
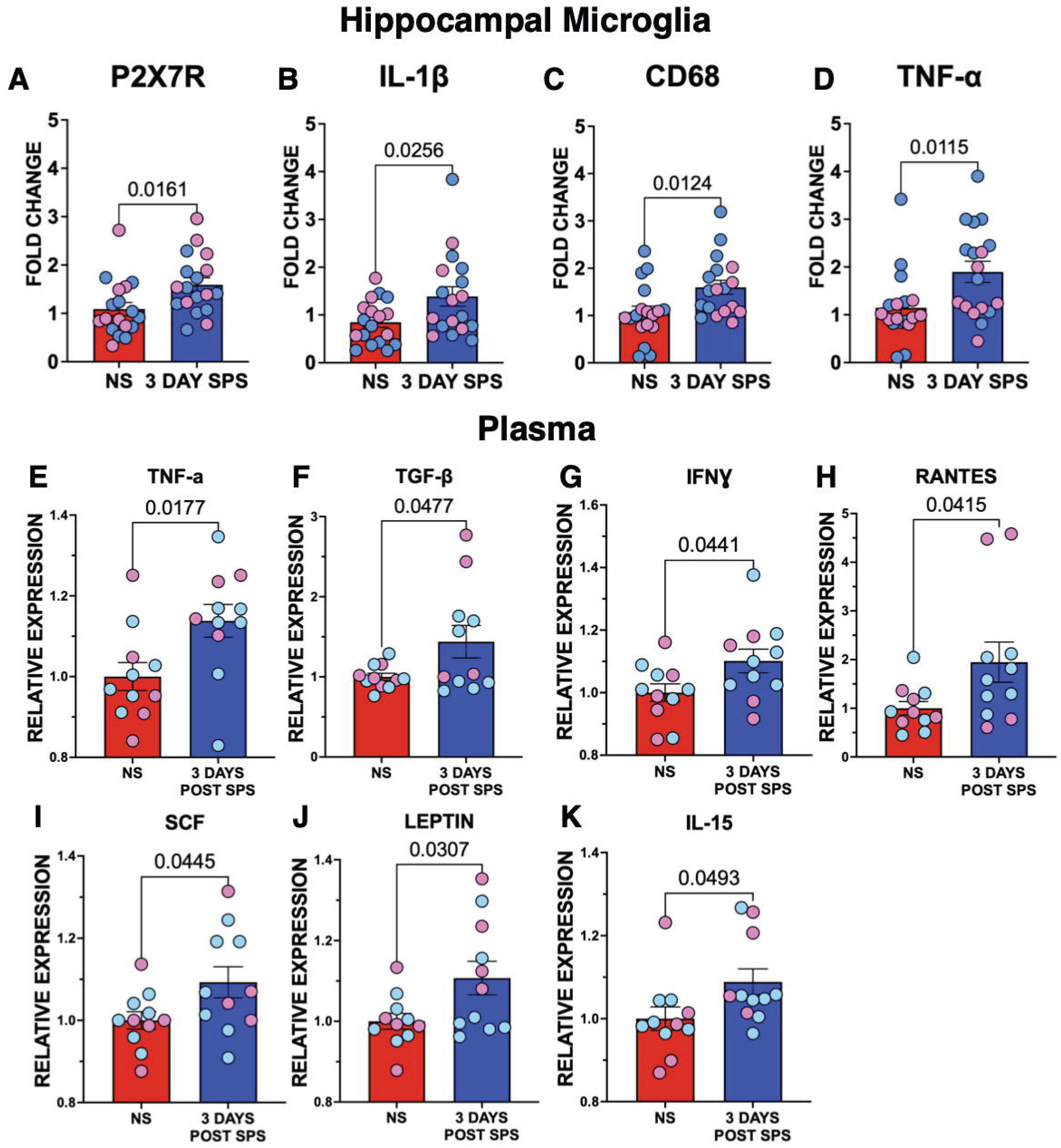
SPS increases hippocampal microglial and peripheral cytokines in male and female rats 3 days after SPS exposure. (A-D) Graphs of the fold change in mRNA expression of hippocampal microglial P2X7R, IL-1β, CD68, and TNF-α genes. Average of duplicate Ct values were normalized to GAPDH expression of the respective male and female NS groups. (E-K) Changes in the peripheral blood cytokines 3 days after SPS exposure. Absorbance values were normalized to respective male and female NS groups. Individual male and female rats shown as light blue and pink circles, respectively.

### Increased peripheral cytokines 3 days post-SPS

To determine whether the SPS-induced inflammatory response also occurred in the blood prior to behavioral training, we measured pro-inflammatory cytokines in serum from male and female rats 3 days after SPS (Figure 3E-K). We found an increase in peripheral blood TNF-α (t (20), 2.586, p = 0.0177), TGF-β (t (20), 2.109, p = 0.0477), IFNƔ (t (20), 2.149, p = 0.0441), RANTES (t (20), 2.178 p = 0.0415), leptin (t (20), 2.326, p = 0.0307), and IL-15 (t (20), 2.093, p = 0.043). No differences were found in the other measured pro-inflammatory cytokines (Supplemental Table 2). Therefore, the SPS-exposed animals entered behavioral training and testing with more inflammatory central and peripheral profiles. Overall, these results suggest that the inflammatory manifestation precedes the observed PTSD-related behaviors in this model.

### P2X7R inhibition prevented the SPS-induced fear extinction impairment and increased anxiety-like behavior

The increased expression of the proinflammatory P2X7R by microglia prior to behavioral training suggests that increased P2X7R signaling could contribute to the PTSD-like phenotype induced by SPS. To test the role of P2X7R signaling in the SPS-associated behavioral changes, we gave ICV infusions of the P2X7R antagonist, A-438079 (3 μg/kg), or vehicle (VEH, 0.9% NaCl Solution) for 7 days starting the day that all animals were exposed to SPS (Figure 4A). As shown in Figure 4B, the VEH-treated and the A-438079-treated male and female rats froze equally to the last CS+ during DAFC on day 2 (t (26), 0.2535, p = 0.8019) and during the CS+ recall on Day 3 (t (26), 0.6122, p = 0.5457), suggesting that P2X7R inhibition did not affect auditory fear acquisition or memory. P2X7R inhibition did produce mild enhancement of fear extinction learning with the A-438079-treated rats showing less freezing at the end of the first EXT session on day 4 (t (182), 2.777, p = 0.0417). In contrast, blocking P2X7Rs had robust effects on extinction recall and anxiety-like behavior (Figure 4C-E). A-438079-treated male and female rats froze less during EXT recall on Day 5 (t (26), 4.498 p = 0.0001) and spent more time in the center of the OFT (t (26), 3.609 p = 0.0013). These results suggest that blocking P2X7Rs prevented the development of PTSD-like behaviors after SPS exposure in both sexes.

**Figure 4:**
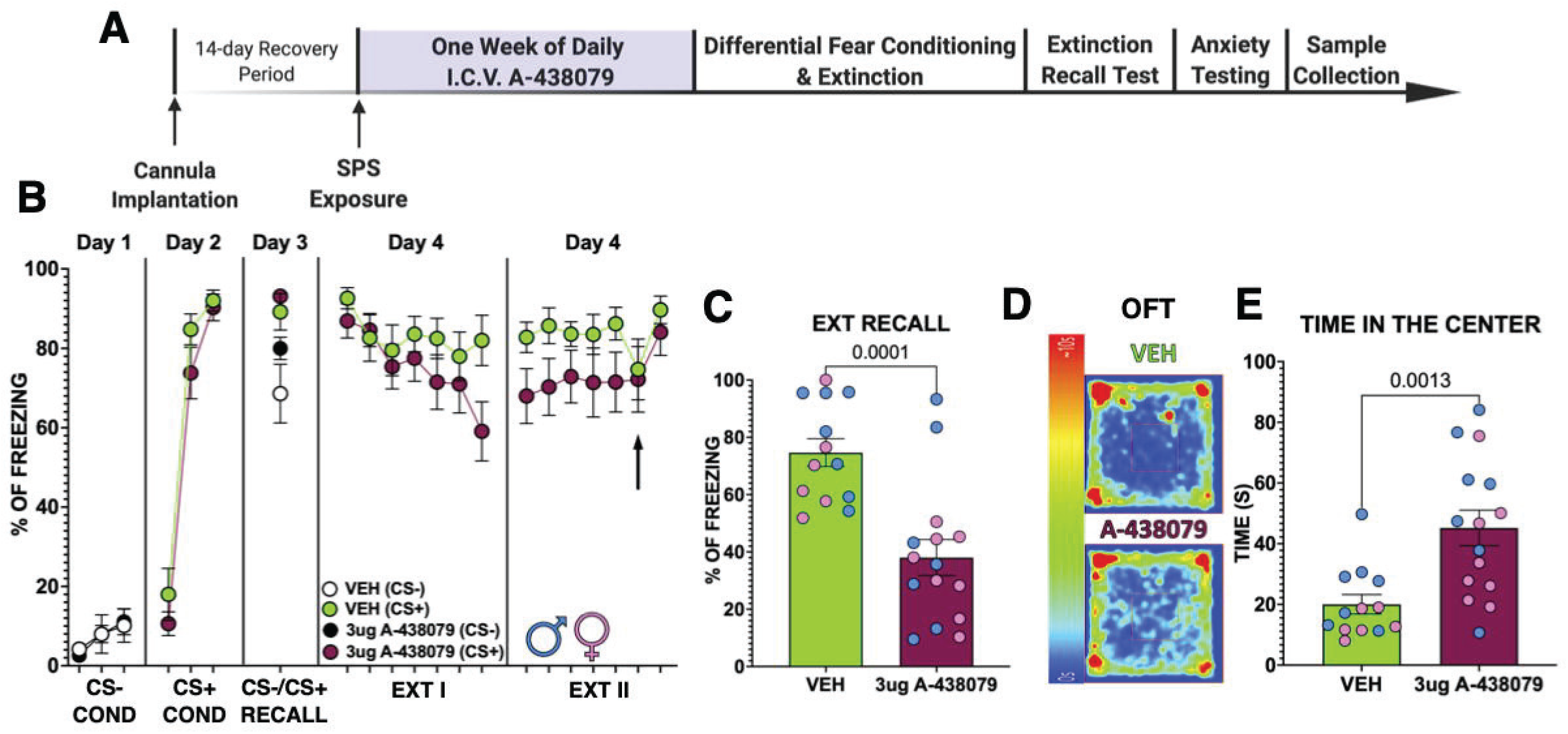
ICV A-438079 prevented SPS from impairing fear extinction recall and increasing anxiety-like behavior in male and female rats. (A) Experimental timeline for the P2X7R blocker experiment. (B) Graph of the percent freezing of male and female rats across the DAFC paradigm. (C) Percent freezing during EXT Recall on Day 5 for male and female rats. (D) Average heat map of center of the body for males and female rats. (E) Graph of the time spent in the center in the OFT for male and female rats. Individual male and female rats respectively shown as light blue and pink circles in C and E.

### Blocking P2X7Rs reduced the expression of Iba1 and P2X7R in the VH

To determine whether blocking P2X7Rs also prevented SPS from increasing the expression of Iba1 and P2X7R in the VH, we immunostained slices obtained after the OFT for Iba1 and P2X7R (Figure 5A-H). Animals infused with A-438079 showed fewer Iba1 (t (20), 2.764 p = 0.0120) and P2X7R (t (20), 3.010 p = 0.0069) positive cells in the VH. In addition, the percentage of area for Iba1 (t (20), 2.165 p = 0.0427) and P2X7R (t (20), 4.740 p = 0.0001) was reduced in A-438079-treated male and female rats.

**Figure 5:**
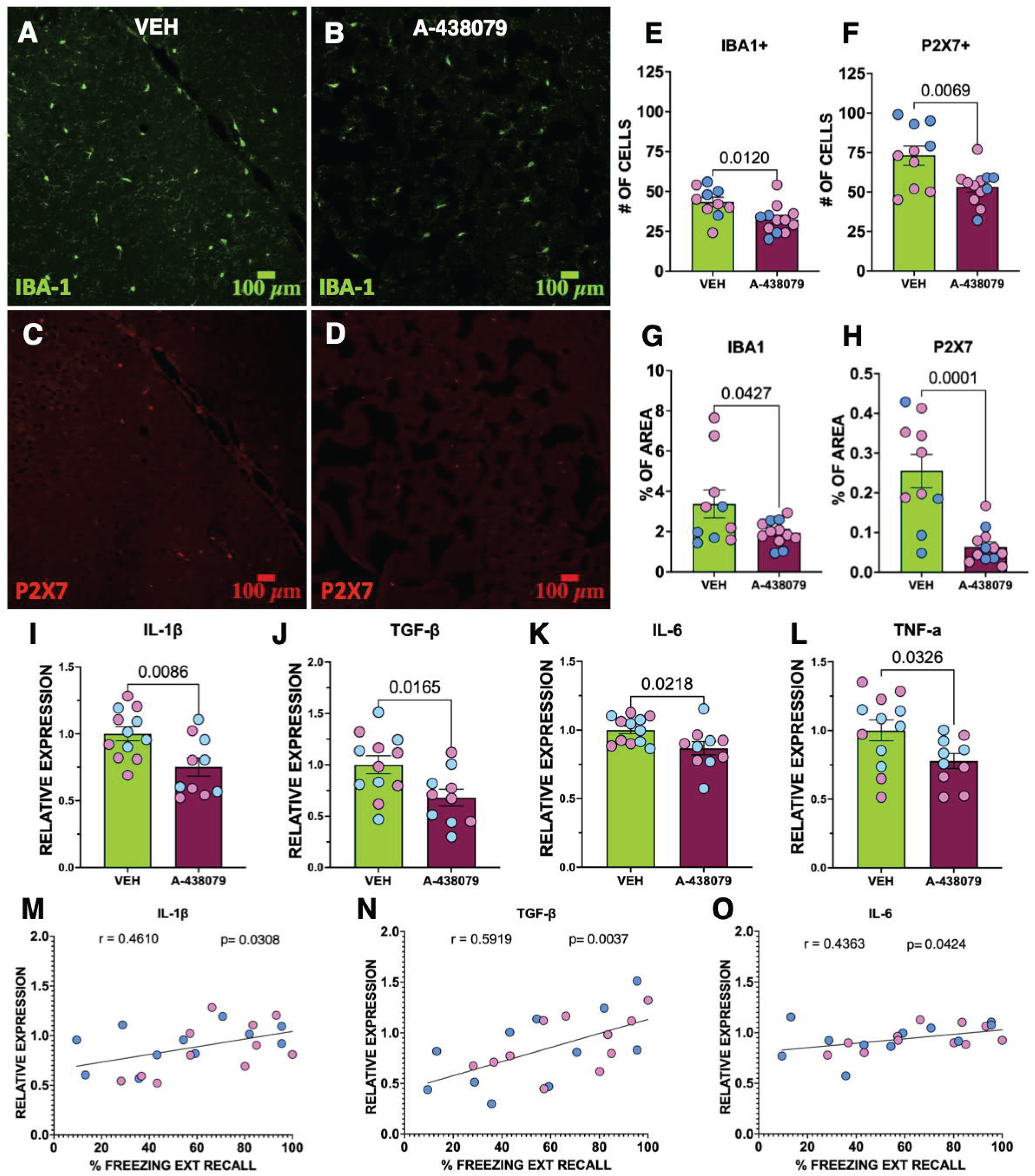
P2X7R Inhibition reduces VH Iba1 and P2X7R expressing cells and peripheral cytokine production in male and female rats. (A, C) Representative images of VH slices co-labeled for Iba1 and P2X7R staining in VEH-treated rats. (B, D) Representative images of A-438079-treated rats. (E-F) Quantification of the number of Iba1+ or P2X7R+ cells in the VH. (G-H) Graphs of the area fraction of Iba1 and P2X7R immunofluorescence in VH slices. (I-L) Changes in the blood cytokines upon P2X7R inhibition. Absorbance values were normalized to the respective male and female VEH groups. (M-O) Correlation of peripheral IL-1β, TGF-β, and IL-6 cytokines and percent freezing during the EXT recall. Individual male and female rats shown as light blue and pink circles, respectively.

### Blocking P2X7Rs reduced peripheral inflammatory cytokines

To determine whether ICV A-438079 affected the peripheral production of cytokines, we measured the protein expression of 16 pro-inflammatory cytokines in plasma collected after the OFT (Figure 5I-L). A-438079-infused male and female animals showed less peripheral IL-1β (t (20), 2.913 p = 0.0086), TGF-β (t (20), 2.617 p = 0.0165), IL-6 (t (20), 2.488 p = 0.0218), and TNF-α (t (20), 2.296 p = 0.0326). No differences were found in other pro-inflammatory cytokines (Supplemental Table 3). Furthermore, the relative expression of Il-1β, TGF-β, and IL-6 positively correlated with the freezing response during EXT recall (Figure 5M-O) indicating that higher cytokine levels were associated with worse extinction recall.

## Discussion

Although the role of the immune system in the pathophysiology of PTSD is unclear, increasing clinical evidence links increased levels of inflammatory cytokines with PTSD [2,26–28]. In our study, we used a well-studied SPS animal model to examine the role of inflammation in the development of PTSD-like behavior after trauma exposure [29–32]. The delayed onset of impaired fear extinction and increased anxiety-like behavior induced by the SPS protocol [7,11,33] allowed us to examine changes in inflammation prior to the onset of the PTSD-like phenotype. We confirmed that SPS exposure leads to impaired fear extinction recall and increased anxiety-like behaviors in males and female rodents one week later. Prior to these behavioral changes, SPS exposure induced higher levels of peripheral and microglial pro-inflammatory cytokines. Furthermore, hippocampal microglia increased expression of the pro-inflammatory P2X7R before the behavioral changes and blocking P2X7Rs prevented the development of the PTSD-like phenotype after SPS exposure, suggesting that stress-induced activation of P2X7Rs and the production of pro-inflammatory cytokines contribute to the development of PTSD-related behaviors.

Consistent with a previous report [7], we also found that SPS-exposed rodents exhibited more microglia in the VH. In addition, more of the microglia expressed P2X7Rs suggesting that SPS increased pro-inflammatory microglia in the VH. Hippocampal microglia showed increased expression of P2X7Rs as soon as three days after SPS exposure which is well before the impaired fear extinction which is manifested one week after SPS [8,11]. At this time point, the microglia also expressed more IL-1β, TNF-α, and CD68 inflammatory genes. This is consistent with reports that SPS increased pro-inflammatory activity in the BLA [7], and increased IL-1β, and TNF-α in the hippocampus [6,34,35]. Our evidence suggests that SPS-induces microglial-mediated inflammation that precedes the impaired fear extinction memory. Fear extinction memory requires VH activity [36,37]. Since elevated IL-1β and TNF-α in the hippocampus can disrupt synaptic plasticity [38] and memory formation [23], their expression by VH microglia could contribute to the impaired fear extinction memory.

In addition to a role in pain disorders [39–41], a growing literature points to the importance of P2X7Rs in stress-induced behavioral changes. Our results suggest that SPS activates P2X7Rs to induce a PTSD-like phenotype in male and female rats. A previous study found that P2X7Rs contribute to chronic restraint stress-induced anhedonia-like and anxiety-like behavior [12]. Furthermore, chronic administration of a P2X7R agonist, BzATP, directly into the hippocampus produced despair-like and anxiety-like behavior in rodents [42]. In addition, the despair-like and anxiety-like behaviors caused by chronic unpredictable stress were prevented by blocking P2X7Rs with A-438079 or using P2X7R KO mice [42]. Thus stress-induced stimulation of P2X7Rs contributes to several behavioral changes which resemble symptoms seen in patients with PTSD.

In addition to increased central inflammation, our data suggest that peripheral inflammation contributes to SPS-induced behavioral changes. SPS-exposed animals displayed an increase in peripheral TNF-α, IFNƔ, TGF-β, IL-15, and RANTES prior to the onset of impaired fear extinction and increased anxiety-like behavior. Also, the animals with impaired fear extinction showed increased peripheral IL-6 and TGF-β. Furthermore, P2X7R blockade reduced peripheral IL-6, TNF-α, IL-1β, and TGF-β and prevented the SPS-induced behavioral changes. These findings are consistent with clinical reports showing increased IL-6, IL-1β, and TNF-α in the blood of patients with PTSD [27,28,43], and suggest that increased peripheral inflammation as a result of a traumatic event might contribute to the development of PTSD perhaps by crossing into the CNS [44].

It is important to note several limitations of our study. First, although we focused on microglial changes in the VH similar inflammatory changes likely occur in other brain structures that regulate fear extinction memory and anxiety-like behavior. Furthermore, the elevated TNF-α and other inflammatory cytokines in blood likely increased inflammation in other brain structures which regulate fear extinction memory and anxiety-like behavior. In addition, since the P2X7R is also expressed by peripheral macrophages and monocytes [45] and astrocytes [46,47] and oligodendrocytes [14] and ICV administration reaches the peripheral circulation [48], we cannot exclude the contribution of P2X7Rs on non-microglial cells in the SPS-induced behaviors.

In conclusion, our findings suggest that the traumatic SPS exposure activates P2X7R signaling to increase inflammation and induce the development of a PTSD-like phenotype of impaired fear extinction and increased anxiety-like behavior. Consistent with the idea that P2X7R expression is associated with mental disorders, higher P2X7R mRNA expression was found in blood samples of treatment-resistant and untreated patients with major depression [49]. Further studies are needed to determine if similar changes in P2X7R expression are seen in patients with PTSD.

## Acknowledgments

This work was supported by the RCMI BRAIN and MAGIC Cores (NIMHHD U54 MD007579), PR-INBRE Institutional Development Award (IDeA) P20GM103475, and R15 MH116345 from the National Institutes of Health. O.T. and Y.R. were supported by the PHSU RISE Program R25-GM082406. A special thanks to Jaime Roman, Anibal Gonzalez Alvarez, Keishla Valentin, Karina Ruiz, and Gabriela Ortiz for their assistance with this project.

## Author contributions

O.T. and J.T.P. designed research; O.T., Y.R., B.V., and M.C. performed research; O.T., Y.R., B.V., M.C., and J.T.P. analyzed data; O.T. and J.T.P. wrote the paper

Keywords: ventral hippocampus, fear extinction, microglia, cytokines, P2X7R, PTSD

