## Supplemental tables for "Purinergic P2X7 Receptor-mediated inflammation precedes PTSD-related Behaviors in Rats"

| Figure | Sample | Statistic | Result |
|---|---|---|---|
| 1B | Males, Day 1, NS (freezing, 5-6 CS-COND) vs SPS (freezing, 5-6 CS-COND) | Unpaired t test, two-tailed | t (26) = 0.5338, p = 0.5980 |
| 1F | Females, Day 1, NS (freezing, 5-6 CS-COND) vs SPS (freezing, 5-6 CS-COND) | Unpaired t test, two-tailed | t (10) = 0.8474, p = 0.3658 |
| 1B | Males, Day 2, NS (freezing, 5-6 CS+COND) vs SPS (freezing, 5-6 CS+COND) | Unpaired t test, two-tailed | t (26) = 0.6821, p = 0.5012 |
| 1F | Females, Day 2, NS (freezing, 5-6 CS+COND) vs SPS (freezing, 5-6 CS+COND) | Unpaired t test, two-tailed | t (10) = 0.7167, p = 0.4899 |
| 1B | NS Males, Day 3, (freezing, CS-RT) vs (freezing, CS+RT) | Unpaired t test, two-tailed | t (26) = 3.436, p = 0.0020 ** |
| 1B | SPS Males, Day 3, (freezing, CS-RT) vs (freezing, CS+RT) | Unpaired t test, two-tailed | t (26) = 3.997, p = 0.0005 *** |
| 1F | NS Females, Day 3, (freezing, CS-RT) vs (freezing, CS+RT) | Unpaired t test, two-tailed | t (10) = 3.613, p = 0.0047 ** |
| 1F | SPS Females, Day 3,(freezing, CS-RT) vs (freezing, CS+RT) | Unpaired t test, two-tailed | t (10) = 2.967, p = 0.0141 * |
| 1B | Males, Day 4, EXT I, NS (CS+ freezing, EXT 13-14) vs SPS (CS+ freezing, EXT 13-14) | 2-way ANOVA, Sidak's Multiple Comparisons | t = 0.7020, DF = 25.25, p = 0.9909 |
| 1F | Females, Day 4, EXT I, NS (CS+ freezing, EXT 13-14) vs SPS (CS+ freezing, EXT 13-14) | 2-way ANOVA, Sidak's Multiple Comparisons | t = 0.4775, DF = 70, p = 0.9991 |
| 1C | Males, Day 5, NS (freezing, CS+EXT-RT) vs SPS (freezing, CS+EXT-RT) | Unpaired t test, two-tailed | t (26) = 4.670, p = 0.0001 *** |
| 1G | Females, Day 5, NS (freezing, CS+EXT-RT) vs SPS (freezing, CS+EXT-RT) | Unpaired t test, two-tailed | t (10) = 2.993, p = 0.0135 * |
| 1E | Males, OFT, NS (Time in the center) vs SPS (Time in the center) | Unpaired t test, two-tailed | t (26) = 2.694, p = 0.0122 * |
| 1I | Females, OFT, NS (Time in the center) vs SPS (Time in the center) | Unpaired t test, two-tailed | t (10) = 2.802, p = 0.0187 * |
| 2E | Males, NS (# of Iba1+ cells) vs SPS (# of Iba1 cells) | Unpaired t test, two-tailed | t (11), 4.064, p = 0.0019 ** |
| 2F | Males, NS (# of P2X7+ cells) vs SPS (# of P2X7+ cells) | Unpaired t test, two-tailed | t (11), 3.129, p = 0.0096 ** |
| 2G | Males, NS (# of Iba1+/P2X7+ cells) vs SPS (# of Iba1+/P2X7+ cells) | Unpaired t test, two-tailed | t (11), 2.998, p = 0.0121 * |
| 2H | Males, NS (Iba1 Area Fraction) vs SPS (Iba1 Area Fraction) | Unpaired t test, two-tailed | t (11) = 1.680, p = 0.1211 |
| 2I | Males, NS (P2X7 Area Fraction) vs SPS (P2X7 Area Fraction) | Unpaired t test, two-tailed | t (10), 2.590, p = 0.0270 * |
| 3A | Males & Females, P2X7R Ct^2 value to NS GAPDH (NS vs 3 days post SPS) | Unpaired t test, two-tailed | t (34), 2.532, p = 0.0161 * |
| 3B | Males & Females, TNFα Ct^2 value to NS GAPDH (NS vs 3 days post SPS) | Unpaired t test, two-tailed | t (34), 2.673, p = 0.0115 * |
| 3C | Males & Females, IL-1β Ct^2 value to NS GAPDH (NS vs 3 days post SPS) | Unpaired t test, two-tailed | t (34), 2.335, p = 0.0256 * |
| 3D | Males & Females, CD68 Ct^2 value to NS GAPDH (NS vs 3 days post SPS) | Unpaired t test, two-tailed | t (34), 2.641, p = 0.0124 * |
| 4B | Males & Females, Day 2, VEH (freezing, 5-6 CS+COND) vs A-438079 (freezing, 5-6 CS+COND) | Unpaired t test, two-tailed | t (26), 0.2535, p = 0.8019 |
| 4B | Males & Females, Day 3, VEH (freezing, CS+ RT) vs A-438079 (freezing,CS+ RT) | Unpaired t test, two-tailed | t (26), 0.6122, p = 0.5457 |
| 4B | Males & Females, Day 4, EXT I, VEH (CS+ freezing, EXT 13-14) vs A-438079 (CS+ freezing, EXT 13-14) | 2-way ANOVA, Sidak's Multiple Comparisons | t = 2.777, DF = 182 p = 0.0417 * |
| 4C | Males & Females, Day 5, VEH (freezing, CS+EXT-RT) vs A-438079 (freezing, CS+EXT-RT) | Unpaired t test, two-tailed | t (26), 4.498 p = 0.0001 *** |
| 4E | Males & Females, OFT, VEH (Time in the center) vs A-438079 (Time in the center) | Unpaired t test, two-tailed | t (26), 3.609 p = 0.0013 ** |
| 5E | Males & Females, VEH (# of Iba1+ cells) vs A-438079 (# of Iba1 cells) | Unpaired t test, two-tailed | t (20), 2.764 p = 0.0120 * |
| 5F | Males & Females, VEH (# of P2X7+ cells) vs A-438079 (# of P2X7+ cells) | Unpaired t test, two-tailed | t (20), 3.010 p = 0.0069 ** |
| 5G | Males & Females, VEH (Iba1 Area Fraction) vs A-438079 (Iba1 Area Fraction) | Unpaired t test, two-tailed | t (20), 2.165 p = 0.0427 * |
| 5H | Males & Females, VEH (P2X7 Area Fraction) vs A-438079 (P2X7 Area Fraction) | Unpaired t test, two-tailed | t (20), 4.740 p = 0.0001 *** |

Statistical Table. Effects of Single Pronlonged Stress on Differential Auditory Fear Conditioning (DAFC), Extinction and Anxiety-like behaviors in male and female Sprague-Dawley rats.

| Proinflammatory cytokines | STATISTICAL ANALYSIS | | | | |
|---|---|---|---|---|---|
| | MEAN | | (B-A) ± SEM | Unpaired (t test) two-tailed | |
| | NON-STRESSED (A) | SPS (B) | | t, df | p value |
| **TGF-β** | 1.000 | 1.257 | 0.2566 ± 0.04927 | t=5.209, df=10 | 0.0004 *** |
| **VEGF** | 1.000 | 1.344 | 0.3438 ± 0.1273 | t=2.701, df=10 | 0.0223 * |
| **IL-6** | 1.000 | 1.530 | 0.5301 ± 0.2248 | t=2.358, df=10 | 0.0401 * |
| **LEPTIN** | 1.000 | 1.156 | 0.1557 ± 0.07527 | t=2.068, df=10 | 0.0655 |
| **IL-1β** | 1.000 | 1.535 | 0.5346 ± 0.2697 | t=1.982, df=10 | 0.0756 |
| **IP-10** | 1.000 | 1.235 | 0.2348 ± 0.1216 | t=1.931, df=10 | 0.0823 |
| **IL-1α** | 1.000 | 1.192 | 0.1923 ± 0.1361 | t=1.412, df=10 | 0.1882 |
| **SCF** | 1.000 | 1.260 | 0.2598 ± 0.1879 | t=1.382, df=10 | 0.1969 |
| **FGFβ** | 1.000 | 1.467 | 0.4672 ± 0.3522 | t=1.326, df=10 | 0.2142 |
| **RANTES** | 1.000 | 1.559 | 0.5592 ± 0.4428 | t=1.263, df=10 | 0.2353 |
| **IL-5** | 1.000 | 2.229 | 1.229 ± 0.9792 | t=1.255, df=10 | 0.2380 |
| **TNFα** | 1.000 | 2.150 | 1.150 ± 0.9297 | t=1.237, df=10 | 0.2444 |
| **IL-15** | 1.000 | 1.663 | 0.6632 ± 0.5378 | t=1.233, df=10 | 0.2457 |
| **IFNɣ** | 1.000 | 2.034 | 1.034 ± 0.8794 | t=1.175, df=10 | 0.2670 |
| **MCP-1** | 1.000 | 1.736 | 0.7358 ± 0.6533 | t=1.126, df=10 | 0.2863 |
| **MIP-1α** | 1.000 | 1.352 | 0.3515 ± 0.3126 | t=1.125, df=10 | 0.2870 |

Supplemental Table 1. Effects of Single Pronlonged Stress on peripheral production of pro-inflammatory cytokines in male rats.

| Proinflammatory cytokines | STATISTICAL ANALYSIS | | | | |
|---|---|---|---|---|---|
| | MEAN | | (B-A) ± SEM | Unpaired (t test) two-tailed | |
| | NON-STRESSED (A) | 3 DAYS POST SPS (B) | | t, df | p value |
| **TGF-β** | 1.000 | 1.439 | 0.4386 ± 0.2079 | t=2.109, df=20 | 0.0477 * |
| **VEGF** | 1.000 | 0.994 | -0.006200 ± 0.05465 | t=0.1135, df=20 | 0.9108 |
| **IL-6** | 1.000 | 0.980 | -0.01999 ± 0.05001 | t=0.3997, df=20 | 0.6936 |
| **LEPTIN** | 1.000 | 1.107 | 0.1071 ± 0.04604 | t=2.326, df=20 | 0.0307 * |
| **IL-1β** | 1.000 | 1.036 | 0.03619 ± 0.05237 | t=0.6912, df=20 | 0.4974 |
| **IP-10** | 1.000 | 1.175 | 0.1752 ± 0.1804 | t=0.9716, df=20 | 0.3429 |
| **IL-1α** | 1.000 | 1.019 | 0.01867 ± 0.06071 | t=0.3076, df=20 | 0.7616 |
| **SCF** | 1.000 | 1.093 | 0.09282 ± 0.04330 | t=2.144, df=20 | 0.0445 * |
| **FGFβ** | 1.000 | 1.083 | 0.08314 ± 0.04862 | t=1.710, df=20 | 0.1027 |
| **RANTES** | 1.000 | 1.948 | 0.9479 ± 0.4352 | t=2.178, df=20 | 0.0415 * |
| **IL-5** | 1.000 | 1.105 | 0.1045 ± 0.06274 | t=1.666, df=20 | 0.1113 |
| **TNFα** | 1.000 | 1.138 | 0.1380 ± 0.05336 | t=2.586, df=20 | 0.0177 * |
| **IL-15** | 1.000 | 1.088 | 0.08844 ± 0.04226 | t=2.093, df=20 | 0.0493 * |
| **IFNɣ** | 1.000 | 1.101 | 0.1014 ± 0.04717 | t=2.149, df=20 | 0.0441 * |
| **MCP-1** | 1.000 | 1.036 | 0.03589 ± 0.05401 | t=0.6646, df=20 | 0.5139 |
| **MIP-1α** | 1.000 | 1.050 | 0.04975 ± 0.04949 | t=1.005, df=20 | 0.3267 |

Supplemental Table 2. Peripheral production of pro-inflammatory cytokines 3 days after SPS exposure in males and female rats.

| Proinflammatory cytokines | STATISTICAL ANALYSIS | | | | |
|---|---|---|---|---|---|
| | MEAN | | (B-A) ± SEM | Unpaired (t test) two-tailed | |
| | VEHICLE (A) | 3µg A-438079 (B) | | t, df | p value |
| TGF-β | 1.000 | 0.6798 | -0.3202 ± 0.1224 | t=2.617, df=20 | 0.0165 * |
| VEGF | 1.000 | 0.9193 | -0.08073 ± 0.08648 | t=0.9335, df=20 | 0.3617 |
| IL-6 | 1.000 | 0.8671 | -0.1329 ± 0.05343 | t=2.488, df=20 | 0.0218 * |
| LEPTIN | 1.000 | 1.1190 | 0.1192 ± 0.2480 | t=0.4807, df=20 | 0.6359 |
| IL-1β | 1.000 | 0.7526 | -0.2474 ± 0.08494 | t=2.913, df=20 | 0.0086 ** |
| IP-10 | 1.000 | 0.9440 | -0.05604 ± 0.05216 | t=1.074, df=20 | 0.2954 |
| IL-1α | 1.000 | 0.8036 | -0.1964 ± 0.1051 | t=1.868, df=20 | 0.0764 |
| SCF | 1.000 | 0.9549 | -0.04508 ± 0.1170 | t=0.3852, df=20 | 0.7041 |
| FGF-β | 1.000 | 0.8340 | -0.1660 ± 0.1059 | t=1.568, df=20 | 0.1326 |
| RANTES | 1.000 | 0.5062 | -0.4938 ± 0.2454 | t=2.013, df=20 | 0.0578 |
| IL-5 | 1.000 | 0.9486 | -0.05140 ± 0.09303 | t=0.5525, df=20 | 0.5868 |
| TNFα | 1.000 | 0.7773 | -0.2227 ± 0.09701 | t=2.296, df=20 | 0.0326 * |
| IL-15 | 1.000 | 0.9670 | -0.03299 ± 0.1633 | t=0.2020, df=20 | 0.8420 |
| IFNγ | 1.000 | 0.7851 | -0.2149 ± 0.1177 | t=1.825, df=20 | 0.0830 |
| MCP-1 | 1.000 | 0.9104 | -0.08965 ± 0.1089 | t=0.8230, df=20 | 0.4202 |
| MIP-1α | 1.000 | 0.8107 | -0.1893 ± 0.1548 | t=1.223, df=20 | 0.2356 |

Supplemental Table 3. Peripheral production of pro-inflammatory cytokines upon ICV administration of a P2X7R antagonist in rats.
